# BSA101: Unlocking Historical Mutant Collections with BSA-Seq

**DOI:** 10.64898/2026.09.24.754120

**Authors:** Brian Zebosi, Comfort Bonney Arku, Madelaine Bartlett, Samuel Leiboff

**Affiliations:** Department of Botany and Plant Pathology, Oregon State University, Corvallis, OR 97331,USA; Department of Biology, University of Massachusetts, Amherst, MA 01003, USA; The Sainsbury Laboratory, University of Cambridge, Cambridge, UK

**Keywords:** BSA-Seq, Forward genetics, Heterogeneous populations, Next-generation sequencing (NGS), Genetic mapping, Maize (Zea mays), *tasselseed*

## Abstract

Forward genetics is a powerful approach for gene discovery, but identifying causal mutations becomes difficult when mutants are maintained in heterogeneous populations with uncertain pedigrees. This is exemplified by classical *tasselseed* (*ts*) mutants, which have long served as a genetic model for studying sex determination and carpel suppression. Decades of repeated outcrossing to diverse inbred lines have created substantial genetic heterogeneity, limiting the effectiveness of conventional bulked-segregant analysis sequencing (BSA-Seq). To address this, we developed a BSA-Seq framework that integrates flexible experimental designs, multiple reference genomes, and complementary statistical methods tailored for genetically heterogeneous populations. Applying this framework revealed that reference genome selection is critical for mapping success and that Euclidean distance raised to the fourth power (ED^4^) outperformed homozygosity mapping (HM). Furthermore, the framework enables simultaneous mapping of multiple mutations within a single population, eliminating the need for additional mapping populations. Applying this framework to 26 *ts* mutant stocks from the Maize Genetics Cooperation Stock Center, we successfully mapped 24 mutants to genomic intervals containing known *ts* genes, while the remaining mutants mapped to distinct genomic intervals, defining novel candidate regions underlying carpel suppression. Together, these results demonstrate that historical mutant collections represent an underutilized resource for gene discovery and establish a generalizable mapping strategy for unlocking their genetic potential across diverse species.

**Author Summary:** Forward genetics has been instrumental in uncovering genes that control important biological processes, but mapping the specific causal mutation responsible for a phenotype can be difficult when mutants are maintained in mixed genetic backgrounds with uncertain pedigrees. This challenge is common in historical mutant collections preserved in public stock centers. Here, we demonstrate that bulked-segregant analysis coupled to whole genome sequencing (BSA-Seq) can effectively map these mutants when combined with flexible experimental designs, multiple reference genomes, and alternative statistical methods. We tested this approach in classical maize *tasselseed* (*ts*) mutants, which alter carpel development and sex determination. Using this approach, we mapped 24 out of 26 mutants to genomic intervals containing known sex determination genes and identified candidate novel intervals for the remaining mutants. Additionally, we showed that multiple independent mutations can be mapped simultaneously from a single population, reducing the time and cost required for genetic mapping. These results demonstrate that historical mutant collections are an untapped resource for gene discovery and provide a clear path for unlocking their genetic potential across diverse species.

## Introduction

How genes contribute to the development of diverse phenotypes remains a fundamental question in biology. Forward genetics has long been one of the most powerful and unbiased approaches for addressing this question by screening mutagenized populations to identify genes underlying phenotypes of interest [1,2]. The mutations exploited in forward genetic screens may arise from natural variation [3] or be experimentally induced through chemical, radiation, or transposon based mutagenesis [4,5], enabling gene discovery across diverse species including zebrafish, mice, Arabidopsis, barley, rice, tomato and maize [6–13]. In maize, transposon systems including *Ac*/*Ds* and *Mutator* have also been widely used to disrupt gene function [14–16]. Radiation mutagenesis, although used less frequently, produces large deletions, insertions, and chromosomal rearrangements such as inversions, duplications, and translocations [17,18]. In contrast, Ethyl methanesulfonate (EMS) remains the predominant mutagen used in maize and induces G/C to A/T nucleotide transitions that generate single nucleotide polymorphisms (SNPs) useful for mutant identification [5].

Once mutants with phenotypes of interest have been identified, the major challenge becomes identifying the causal mutation, which can be accomplished through linkage mapping using individual segregant analysis (ISA) or bulked segregant analysis (BSA) coupled with positional cloning [19,20]. ISA requires genotyping large numbers of individual segregants, making it labor-intensive [20], whereas BSA simplified this process by pooling segregants with contrasting phenotypes into separate bulks for sequencing to identify genomic regions linked to the causal mutation [20–23]. More recent advances in next-generation sequencing and reduced costs have further transformed this approach into BSA coupled with whole-genome sequencing (BSA-Seq), enabling rapid mapping of causal mutations to small genomic intervals [23,24].

Despite its effectiveness, most existing BSA-Seq frameworks have been developed and optimized for mutants derived from uniform genetic backgrounds with known parental pedigrees. Under these conditions, reference genome selection is straightforward, and commonly used statistical methods such as homozygosity mapping perform predictably [23]. However, as BSA-Seq expands to complex populations and non-traditional model species, these assumptions may not hold [25–28], and clear guidance for selecting appropriate reference genomes and statistical methods is lacking. This challenge is exemplified in maize, where many mutants maintained in stock collections or exchanged among research groups have been repeatedly outcrossed to diverse inbred lines, resulting in heterogeneous genetic backgrounds with uncertain pedigrees [29]. Although recent allelism tests of dwarf mutants from the Maize Genetics Cooperation Stock Center demonstrate that these collections harbor extensive allelic diversity at known loci [30], robust strategies for mapping causal mutations in such populations remain limited.

Here, we present a BSA-Seq framework designed to accommodate uncertainty in parental pedigree and reference genome selection. We apply this framework to map 26 *tasselseed* (*ts*) mutants obtained from the Maize Genetics Cooperation Stock Center [31]. Tasselseed mutants are an extensively studied class of developmental mutants in maize that disrupt sex determination and carpel suppression, resulting in the persistence of pistils and kernel production in otherwise staminate tassels [32–40]. Recovered from historical mutagenesis screens and subsequently outcrossed to diverse inbreds, these mutants segregate in heterogeneous genetic backgrounds, making them an ideal system for evaluating our framework. Ultimately, this work provides a generalized BSA-Seq strategy for complex populations while expanding the genetic resources available for floral development and sex determination.

## Results

### BSA-Seq experimental designs and mapping pipeline

To map 26 *ts* mutants that segregated in heterogeneous genetic backgrounds with uncertain pedigrees, we evaluated three BSA-Seq experimental mapping designs across F2 populations (**Fig 1A**) and assessed how reference genome selection influenced mapping performance. In the first design, a mutant-only approach [23,41], only mutant individuals were pooled and sequenced. While cost-effective, the design lacked a wild-type pool to account for background variation and relied entirely on the reference genome to localize causal mutations **(Fig 1B)**. The second design was a classical wild-type versus mutant design [19], in which separate pools of wild-type and mutant individuals were sequenced and allele frequency difference (AFD) used to identify genomic regions linked to the causal mutation **(Fig 1C)**. Finally, for populations segregating for more than one mutant phenotype, we employed a multi-mutant design that enabled simultaneous mapping of multiple mutants within a single population, eliminating the need to generate separate mapping populations for each mutant **(Fig 1D)**.

**Fig 1.**
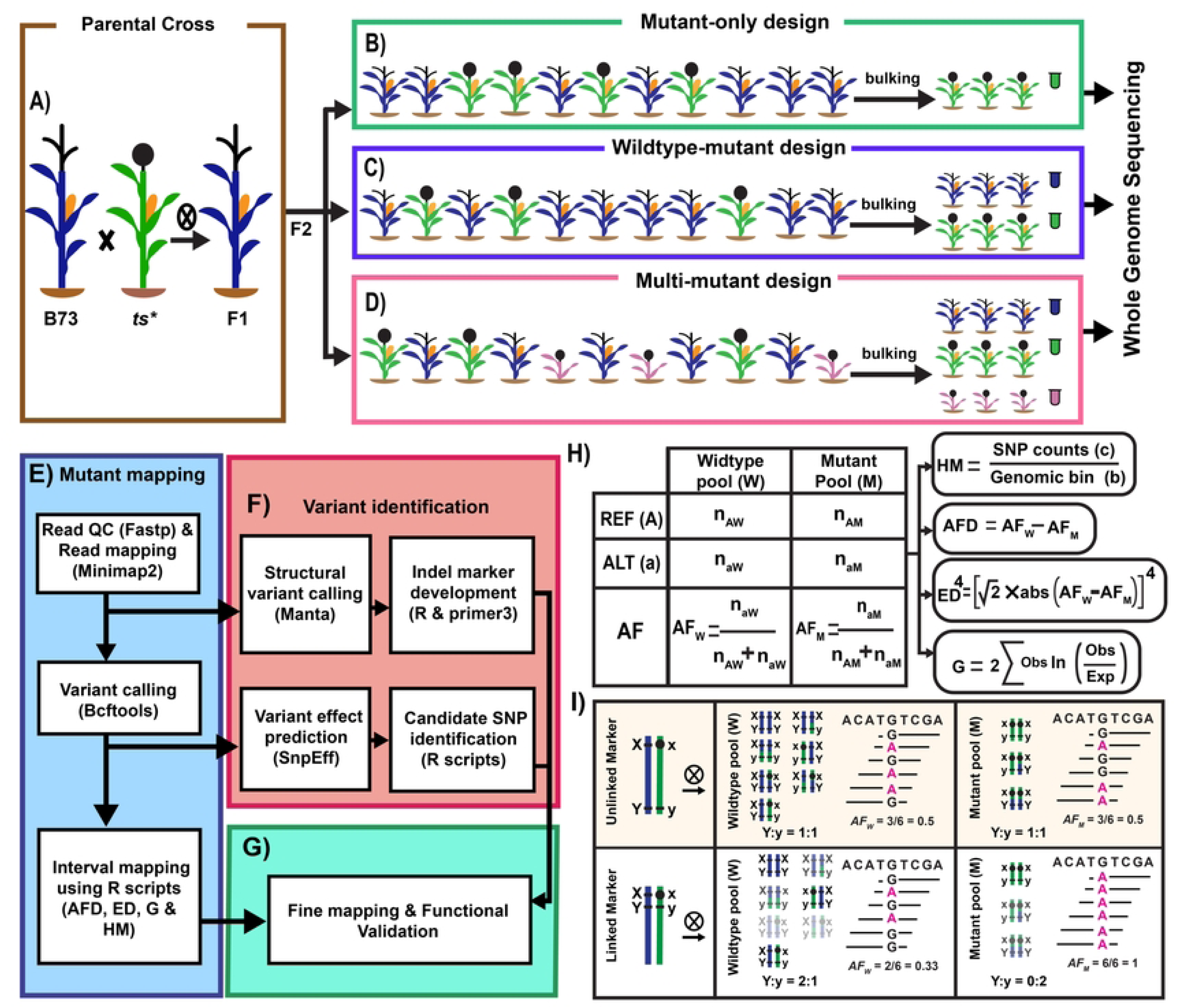
Experimental design and BSA-Seq bioinformatic pipeline for mapping mutants in maize. **A-D)** F2 mapping populations were analyzed using three BSA designs (mutant-only, wild-type vs mutant and multi-mutant), and DNA bulks subjected to whole-genome sequencing. **E-G)** The BSA-Seq pipeline comprises read quality control, read alignment, variant calling, interval mapping using allele frequency difference (AFD), Euclidean distance raised to the fourth power (ED^4^), G-statistic (G), and homozygosity mapping (HM), followed by identification of structural variants and candidate SNPs. (**H)** Allele frequencies (AFs) derived from reference (A) and alternate (a) read counts in wild-type (W) and mutant (M) bulks were used to calculate AFD, ED^4^, HM and G (where Obs and Exp denote observed and expected allele counts, respectively). **I)** Expected AFs for unlinked and linked SNP markers. Blue and green chromosomes denote the parental haplotypes, with mixed segments indicating recombination. The causal locus carries the wild-type (X) and mutant (x) alleles, whereas Y and y denote codominant marker alleles distinguishing B73 and mutant parents. Unlinked markers assort independently, with similar AFs between bulks, while tightly linked markers show limited recombination, yielding skewed AFs and co-segregation with the mutant phenotype.

Across all designs, sequencing reads were aligned to five maize reference genomes including B73, Oh43, P39, Mo17, and W22 were selected based on available pedigree information and to represent genetic diversity [42–44], and variants were called using a customized bioinformatic pipeline (**Fig 1E-G**). Allele frequencies (AFs) were calculated for each SNP from reference and alternate read counts, and additional statistical methods, including allele frequency difference (AFD) [45], G-statistics (G) [46], homozygosity mapping (HM) [23] and Euclidean distance raised to the fourth power (ED^4^) [47], were applied to map genomic regions linked to causal mutations and define mapping intervals (**Fig 1H**). In BSA-Seq, SNPs unlinked to the causal mutation are expected to segregate at a 1:1 ratio, resulting in AFs close to 0.5 without a distinct mapping peak [45]. Conversely, SNPs tightly linked to the causal mutation are expected to co-segregate with the mutant phenotype, yielding mutant AFs approaching 1 with a localized mapping peak that defines the candidate interval [45]. Candidate intervals were subsequently examined for co-segregating structural variants and high confidence SNPs. The functional impacts of prioritized SNPs were predicted using snpEff [48], and molecular markers were designed from prioritized SNPs and structural variants for fine mapping and functional validation (**Fig 1F-G**). Using this integrated approach, we next evaluated how reference genome choice influenced mapping performance across experimental designs.

### Reference genome selection determines mapping success in mutant-only BSA-seq

To evaluate how reference genome choice influences mapping performance in the mutant-only BSA-Seq design, we mapped the *ts*-N2490* mutant (*4012U*), which segregated in a heterogeneous genetic background. Sequencing reads from the mutant pool were independently aligned to four reference genomes (B73, Mo17, Oh43, and P39), and mapping performance was assessed using AF and HM. Alignment to B73 revealed a clear mapping peak on chromosome 1 **(Fig 2A)**, while alignments to Mo17, Oh43, and P39 failed to yield a distinct mapping signal with either AF or HM due to increased background noise that obscured the true peak (**Fig 2B-D**).

**Fig 2.**
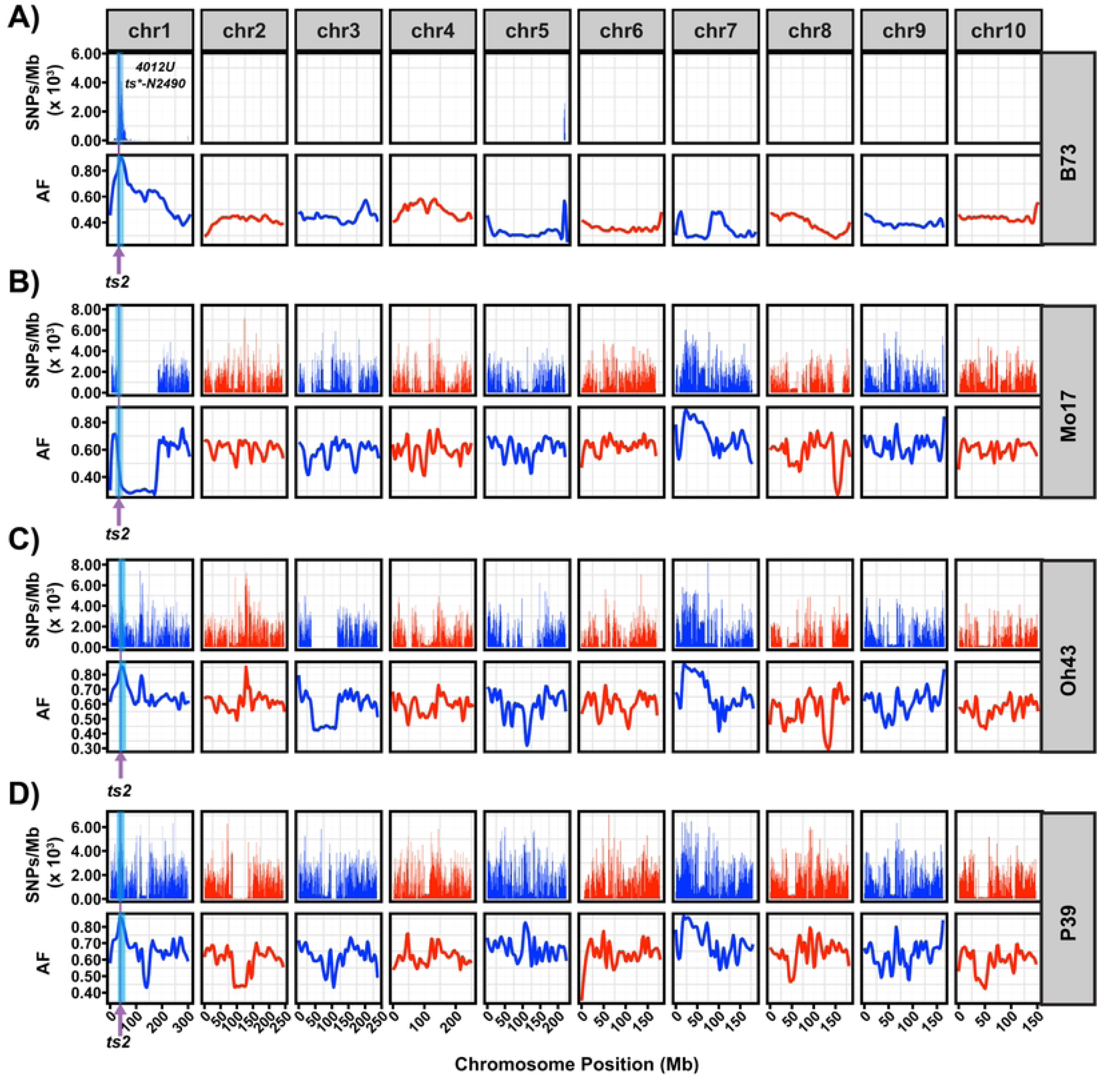
Mutant-only BSA-Seq mapping depends on reference genome selection. **A-D)** Reads from the 4012U (*ts*-N2490*) mutant pool aligned to four reference genomes (B73, Mo17, Oh43 and P39), showing genome-wide Homozygous SNP counts (AF ≥ 0.99) per 1 Mb and allele frequency (AF). **A**) Alignment to B73 localized the mutation to a ∼25–55 Mb interval on chromosome 1 (light blue shaded region), containing the *ts2* locus (∼46.5 Mb, purple arrow). **B-D)** In contrast, alignments to Mo17, Oh43 and P39 exhibited extensive background homozygosity and diffuse AF patterns without distinct mapping peaks.

The differences in mapping performance across alignments likely reflect differences in genetic relatedness between each reference genomes and the mutant background. The distinct mapping peak obtained with B73 is consistent with prior backcrossing to B73, which is expected to reduce background polymorphism [41]. In contrast, alignments to Mo17, Oh43 and P39 resulted in extensive background polymorphism, making the causal mapping region difficult to distinguish from the genetic background. Many of the homozygous variants observed with these reference genomes likely represent genetic differences between the mutant background and each reference genome rather than variants associated with the causal mutation. Consequently, homozygosity mapping becomes less informative as background homozygosity increases. The successful mapping of *4012U ts*-N2490* using B73 is consistent with previous findings indicating that homozygosity mapping and mutant-only BSA-Seq perform best when the reference genome closely resembles the genetic background of the mutant [23,41,49,50].

Overall, these results demonstrate that successful mutant-only mapping depends heavily on reference genome selection.

### ED^4^ outperforms homozygosity mapping across reference genomes

Given that reference genome choice greatly impacted the success of homozygosity mapping (HM), we next evaluated whether alternative statistical approaches could provide more consistent mapping results in heterogeneous genetic backgrounds when different reference genomes are used. We compared HM with several commonly used comparative statistical approaches, including AFD, ED^4^, and G. We assessed the performance of these methods using a classical wild-type versus mutant design across multiple reference genomes. Among the comparative methods tested, ED^4^ produced the highest mapping resolution, defined as the narrowest mapping interval spanning the causal mutation [51]. Compared with AFD and G, ED^4^ generated narrower intervals with reduced background noise (**S1 Fig**), consistent with previous reports that ED^4^ amplifies true mapping signals while suppressing background noise [47]. We therefore selected ED^4^ for subsequent comparisons with HM.

To evaluate the robustness of ED^4^ and HM across genomes, we mapped *ts*-04HI* (6609A ), which segregated in a heterogeneous genetic background (6609A ts*-04HI-A632xOh43 GN-36/B73). Sequencing reads from the mutant and wild-type pools were aligned to four genomes (B73, Mo17, Oh43 and P39). Read alignment rates were uniformly high across all genomes (97.9-98.9%), yielding mean genome coverages of 22x and 27x for wild-type and mutant pools, respectively (**S2 and S3 Tables**). Across all four genomes, ED^4^ consistently recovered a clear mapping peak on chromosome 2. In contrast, HM performance was strongly dependent on reference genome choice, yielding a distinct region of enriched homozygosity on chromosome 2 only when reads were aligned to B73 (**Fig 3A**), whereas alignments to the other three genomes exhibited extensive genome-wide homozygosity that obscured the true mapping signal (**Fig 3B-D)**. These results demonstrate that ED^4^ is more robust than HM in genetically heterogeneous populations across different reference genomes.

**Fig 3.**
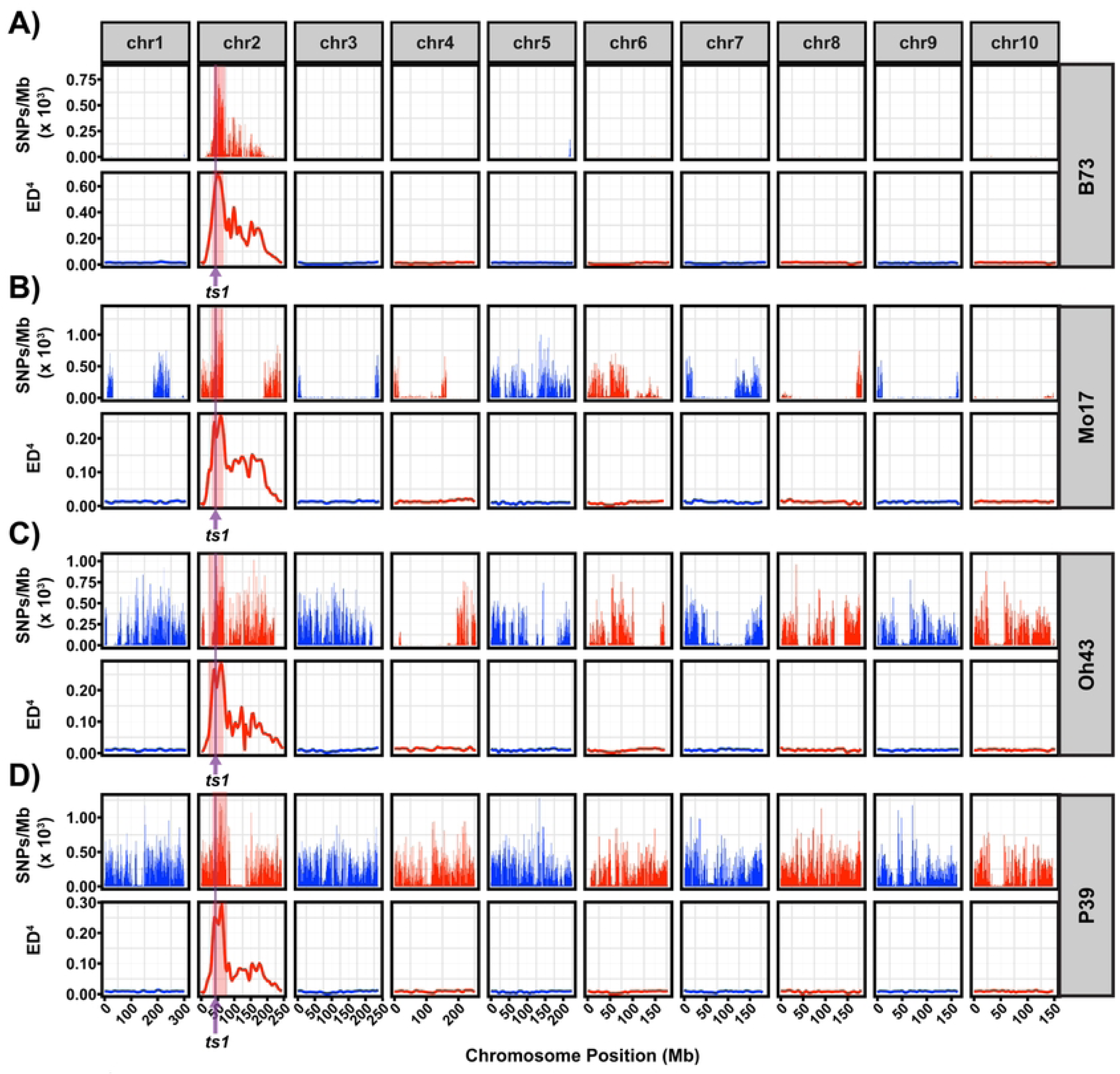
ED⁴ is more robust to reference genome selection than homozygosity mapping. **A-D)** BSA-Seq mapping of *6609A* (*ts*-04HI*) mutant using a wild-type versus mutant design, with reads aligned to B73, Mo17, Oh43, and P39. **A)** Alignment to B73 produced a distinct homozygosity peak on chromosome 2, mirrored by a strong ED⁴ signal. The interval spanned 41.6-49.9 Mb (light red shaded region) and contained the *ts1* locus (∼47 Mb ; arrow). **B-D).** Conversely, alignments to Mo17, Oh43, and P39 failed to yield a homozygosity peak, whereas ED⁴ consistently identified a distinct peak on chromosome 2.

### Multi-mutant design enables mapping of multiple mutants in a single F2 population

To assess whether multiple mutants could be mapped simultaneously from a single F2 population, we analyzed the 6508K *ts*-07IL* population, which segregated two non-allelic tasselseed mutants: a normal-height tasselseed mutant (*ts\**) and a dwarf *tasselseed* mutant, hereafter designated *tiny tasselseed* (*tts*-2481*). We applied a multi-mutant design by sequencing three phenotypically distinct pools (22 plants per pool) representing wild-type, *ts\** and *tts\** individuals. Read alignment rates to the B73 reference genome were 98.1%, 98.9% and 97.7% for wild-type, *ts\** and *tts\** pools, respectively, yielding mean genome coverages of 22.8x, 26.2x and 21.4x (**S2 and S3 Tables**).

To localize the causal mutations underlying *ts\** and *tts\** phenotypes, we used HM and ED⁴ across three pairwise pool comparisons (wild-type versus *ts*^◻^, wild-type versus *tts*^◻^, and *ts*^◻^ versus *tts*^◻^). HM identified regions of enriched homozygosity on chromosome 2 for *ts\** and chromosome 1 for *tts\** (**Fig 4A**). Similarly, ED^4^ identified the same mapping peaks in both wild-type comparisons (**Fig 4B**). Notably, direct comparison of the *ts*^◻^ and *tts*^◻^ mutant pools recovered both mapping peaks without a wild-type pool (**Fig 4C**). These results demonstrate that a multi-mutant design enables simultaneous mapping of multiple mutations in a single population without generating independent mapping populations.

**Fig 4.**
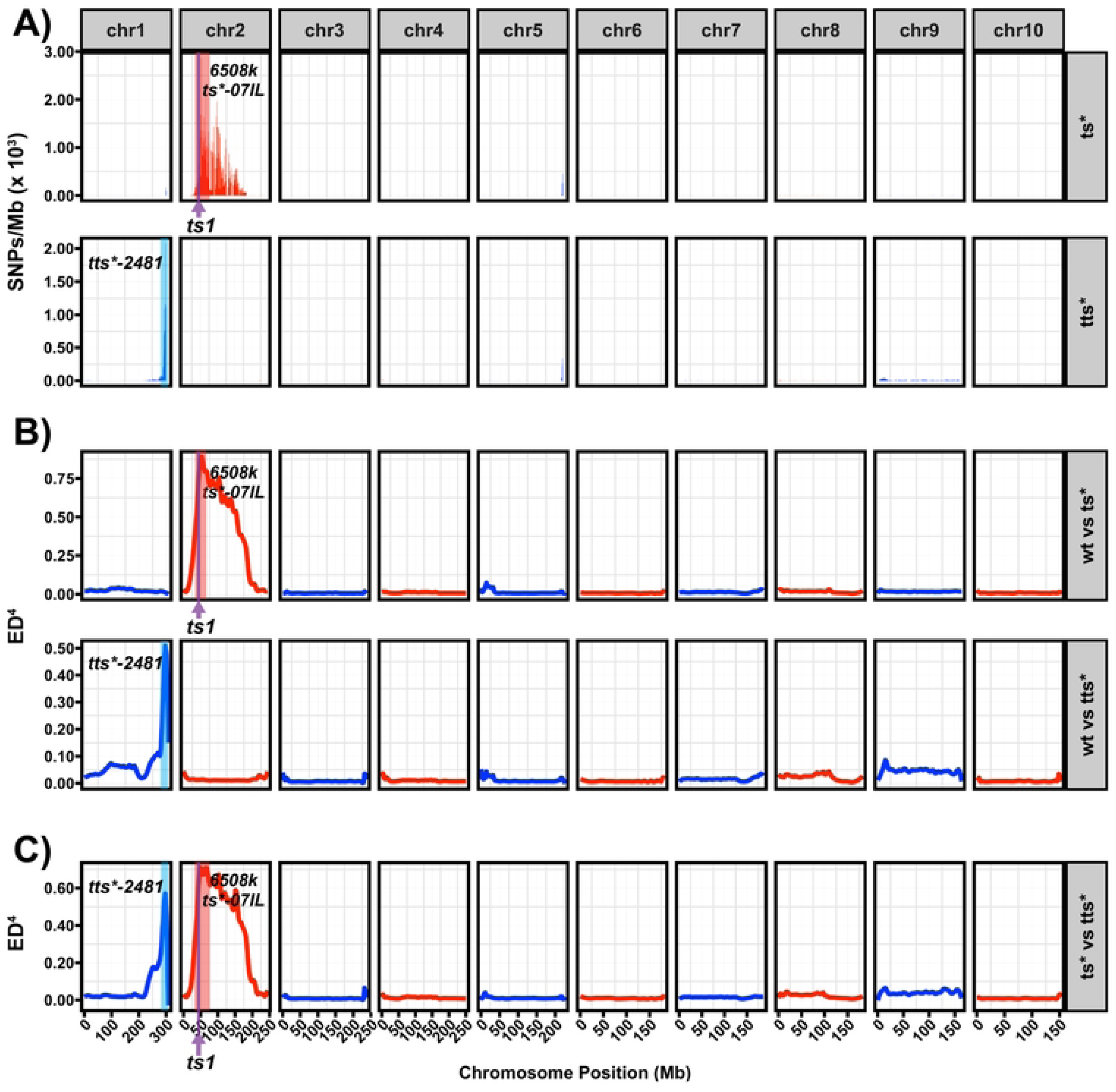
Multi-mutant BSA-Seq design simultaneously maps two mutants using a single mapping population. **A-C).** Reads from wild-type (wt), normal-height *tasselseed* (*ts*-07IL*) and *tiny tasselseed* (*tts*-2481*) mutant pools were aligned to B73. **A).** Homozygous SNP counts (AF ≥ 0.99) per 1 Mb for *ts*-07IL* (top) and *tts*-2481* (bottom). **B)**. ED⁴ for wt versus *ts*-07IL* (top) and wt versus *tts*-2481* (bottom). **C)** ED⁴ comparison between the *ts*-07IL* and *tts*-2481* pools. Both HM and ED⁴ localized the *ts*-07IL* mutation to a 7.4 Mb interval on chromosome 2 (46.4– 53.8 Mb, light red shaded region) containing the *ts1* gene (∼47 Mb, arrow), whereas *tts*-2481* mapped to a 5.1 Mb interval on chromosome 1 (294–299 Mb, light blue shaded region). Direct comparison between the *ts*-07IL and tts*-2481* pools identified both mapping intervals.

### Most *tasselseed* mutants map to genomic regions containing known *tasselseed* loci

Using our BSA-Seq mapping framework, we mapped 26 unmapped tasselseed mutants from the Maize Genetics Cooperation Stock Center and found that 24 mapped to genomic regions harboring known *tasselseed* genes including *tasselseed1* (*ts1*), *tasselseed2* (*ts2*), *tasselseed4* (*ts4*), *Tasselseed 6* (*Ts6*) and *VRS1-like1* (*VRL1*) [32,33,38,39,52]. Specifically, the mapping intervals included the *ts2* locus on chromosome 1 (7 mutants), *ts1* locus on chromosome 2 (14 mutants), the *Ts6* , *ts4* and *VRL1* loci on chromosomes 1, 3 and 7, respectively (1 mutant each; **Fig 5, S2, S3 and S4 Figs**). The localization of these mutants to intervals containing known *tasselseed* loci suggests that many of these mutants are novel alleles of these genes. The remaining two mutants, *tts*-2481* and *ts*-N1967A* mapped to regions on chromosomes 1 and 4, respectively, that contained no known *tasselseed* genes, suggesting that they represent novel loci (**Fig 5; S4 Table**).

**Fig 5.**
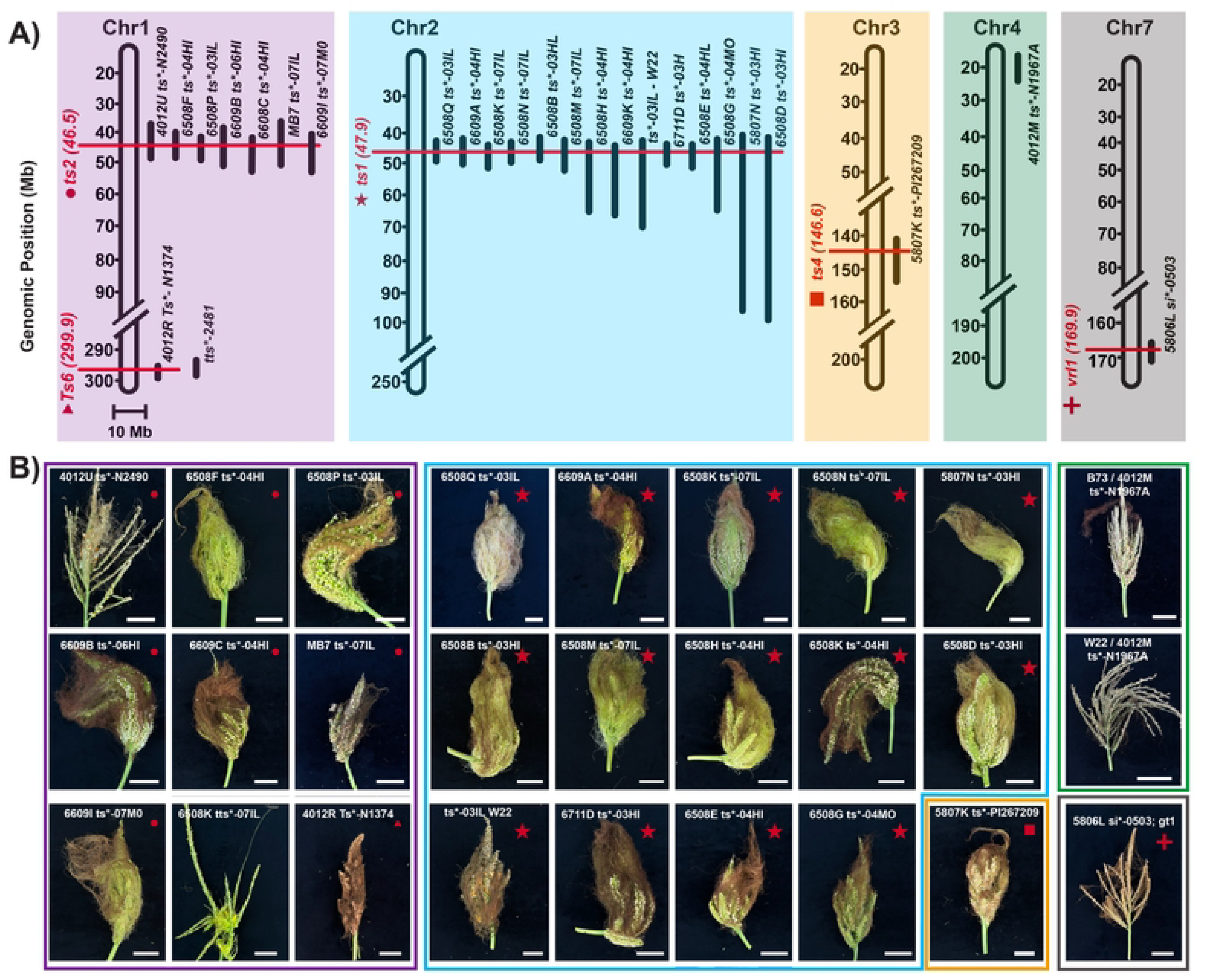
Mapping intervals and phenotypes of 26 *tasselseed* mutants. **A)** Genomic mapping intervals of the 26 tasselseed mutants across chromosomes 1, 2, 3, 4 and 7. Vertical black bars represent mapping intervals with length reflecting relative interval sizes, and red horizontal lines indicate the genomic positions of the known *tasselseed* genes (*Ts1*, *Ts2*, *Ts6*, *Ts4* and *Vrl1)*. Intersections between mapping intervals and gene positions indicate that the corresponding gene resides within the mapped interval. **B)** Tassel phenotypes of the mapped tasselseed mutants. Colored image borders correspond to the chromosome on which each mutant mapped (matching panel A), and red shapes in the upper right of each image correspond to known *tasselseed* genes indicated in panel A. Scale bars = 5 cm.

To identify candidate causal variants, we filtered for canonical EMS induced nucleotide transitions (G/C to A/T) and removed known maize HapMap variants, prioritizing variants by their predicted effects on gene function. Among the seven mutants whose intervals contained *ts2*, three carried missense mutations in *ts2*. Similarly, 5 of the 14 mutants with intervals spanning *ts1*, harbored moderate or high impact variants in *ts1* (**S4 Table**). Notably, the remaining mutants lacked predicted coding mutations at either *ts1* or *ts2*. These findings raise the possibility that some carry variants in cis-regulatory regions of these genes although additional fine mapping and functional validation will be required to identify the causal mutations.

Among the mutants mapping to novel genomic intervals, *4012M* (*ts*-N1967A*) localized to a 15-Mb interval spanning 10–25 Mb on chromosome 4 (**Fig 5A, S2 and S5 Fig**). Because the mutant exhibited weaker expressivity in the W22 background than in B73 **(Fig 5B)**, we generated independent mapping populations in both backgrounds. Despite the reduced expressivity in W22, both populations mapped the mutation to the same genomic interval (**S5 Fig**). Notably, the interval contained no known *tasselseed* genes or predicted functional variants, suggesting that *ts*-N1967A* represents a novel *tasselseed* locus, although additional mapping will be required to identify the causal lesion.

The second mutant mapping to a novel interval, *tiny tasselseed* (*tts*-2481*), arose spontaneously in the 6508K tasselseed stock, exhibited dwarfism, and segregated as a single recessive mutation (χ² = 1.25, df = 1, P = 0.264). About 40% of these mutants were also tasselseed, although whether this results from *tts*-2481* or a genetic interaction with the original 6508K mutation remains unclear. On chromosome 1, *tts\** mapped to a 290-300Mb interval (**Figs 4 and 5, S2 Fig**), containing one splice acceptor SNP and six missense SNPs in genes encoding a nucleotide-binding site-leucine-rich repeat protein, polygalacturonases, C2C2-DOF transcription factor 27 (*dof27*), and two proteins of unknown function (**S4 Table**). Among these genes, *Zm00001eb062640* (*dof27*) represents the top candidate because it contains a splice acceptor variant predicted to disrupt transcript splicing, and DOF transcription factors have established roles in cell proliferation, vascular development, and floral organ development [53,54].

Among the mutants mapping to intervals harboring characterized *tasselseed* loci, *4012R* (*Ts*-N1374*) segregated as a semi-dominant mutation and showed dosage-dependent phenotypic expressivity. Among 40 progeny, 15 exhibited a normal tassel phenotype, whereas 16 and 9 plants displayed weak and strong mutant phenotypes, respectively, consistent with a 1:2:1 segregation ratio (χ² = 2.15, df = 2, P = 0.341). *4012R* mapped to a 290-300Mb genomic region on chromosome 1 containing *Zm00001eb062460* (*Ts6*), but no variants predicted to disrupt *Ts6* function were identified (**Fig 5, S4 Table**). However, the inheritance pattern and phenotype of *4012R* closely resembled those of *Ts6* mutants [52], supporting *Ts6* as the primary candidate gene, although the causal mutation remains to be determined.

The recessive mutant *5806L* (*si*-0503*) displayed no obvious phenotype on its own but produced feminized tassels with long silks in a *gt1* background [37]. The mutation mapped to a 2.7 Mb interval on the long arm of chromosome 7, where no predicted functional variants were identified (**Fig 5, S2 Fig, S4 Table**). The interval included *Zm00001eb325630* (*ZmHB13/VRL1*), encoding a homeodomain-leucine zipper (HD-ZIP) transcription factor implicated in carpel suppression [34,37–40]. Because the *si*-0503;gt1* double mutant phenocopies *gt1;vrl1* double mutants [38,39], *VRL1* represents a candidate gene for *si*-0503*.

Mutant *5807K* (*ts*-PI267209*) segregated as a recessive mutation (χ² = 0.8, df = 1, P = 0.371) and produced highly branched tassels densely covered with long silks (**Fig 5**). Using BSA-Seq, *5807K* mapped to a 2.8 Mb interval on chromosome 3 (**Fig 5**, **S2 Fig**), that lacked functional predicted variants but contained *tasselseed4* (*ts4*), whose mutant phenotype resembled that of *5807K [52]*. Because *ts4* is unannotated in the B73v5 genome assembly, we aligned the B73v3 *ts4/zma-MIR172e* sequence to B73v5 to identify the corresponding region. Two SNPs were identified within the *ts4* locus at chr3:146,624,466 (C>T) and chr3:146,625,041 (C>T) (**S4 Table**), neither of which were located in the mature miR172 sequence. However, one was located in the 5’ region of the pri-microRNA, near the promoter region harboring Helitron insertions responsible for the *ts4* mutant phenotype [52]. These observations support *ts4* as a likely candidate gene underlying the 5807K phenotype

## Discussion

Identifying the genetic basis of mutant phenotypes remains a central goal of forward genetics, providing unbiased insights into molecular mechanisms underlying biological processes [1,2]. Bulked segregant analysis combined with whole genome sequencing (BSA-Seq) has enabled rapid mapping of mutants in maize and other species [23,27,55]. However, most applications have focused on mutants in genetically uniform backgrounds derived from controlled mapping populations and analyzed relative to single reference genomes. While this can be successful for lab-generated mutants, many mutants in stock centers or derived from horticulture arise spontaneously, or segregate in heterogeneous genetic backgrounds with unknown pedigree information [29,36,56–62]. Heterogeneous backgrounds often contain standing genetic variation that can reduce mapping resolution and obscure causal mutations [47]. Homozygosity mapping, in particular, relies on local fixation around causal mutations, an assumption that is often violated in heterogeneous populations because genetic diversity masks mapping signals by generating spurious homozygosity peaks unrelated to causal loci [23,63]. Here, we present a framework that includes experimental design, reference genome choice, and statistical metrics to enable mapping in genetically complex populations. By explicitly accounting for genetic diversity, this framework extends BSA-Seq beyond uniform genetic backgrounds, including to diverse mutant collections from genetically complex populations.

Using this framework, we mapped 26 stock center mutants and identified candidate regions for 24 mutants overlapping intervals for previous cloned sex determination genes, including two loci not previously associated with carpel suppression and sex determination in maize. These findings highlight the value of genetic resources preserved at Maize Genetics Cooperation Stock Center, which maintains decades of rare mutants that would be lost or difficult to recreate with modern approaches, yet thousands of mutants remain unmapped and uncharacterized [64]. In addition, these collections provide a series of alleles of mutants spanning a range of phenotypic expressivity, offering opportunities to dissect gene function and genetic interactions [65,66]. However, public stock centers also present challenges, as many mutants have undergone historical outcrossing or have unknown mutation origins, complicating mapping efforts [29,59]. Importantly, advances in genomics and sequencing technologies, as implemented in our framework, make it possible to revisit previously intractable mutants and identify the genetic basis of their phenotypes.

We demonstrate that BSA-Seq enables cost-effective simultaneous mapping of multiple mutants and genetic modifiers in a single population. In public seed stocks, uncontrolled outcrossing to diverse inbred genetic backgrounds introduces additional segregating variants, including spontaneous mutations that can modify mutant phenotypes and complicate mapping efforts [29,67]. More broadly, this design could be useful for cost-effective mapping in populations segregating multiple independent mutants, or for mapping double or other higher-order mutants. Successful implementation of this would require that individual mutant phenotypes be clearly distinguishable to ensure accurate bulking and minimize cross contamination between pools as mutants of similar phenotype would make phenotyping of mapping population challenging. For example, a mapping population segregating with 2-4 distinct recessive mutants, pool size of 15-30 individuals would be sufficient to localize causal lesions [23], that way sequencing depth can be increased by shifting resources from wild-type sibling pools. Rather than treating segregating variants in stocks obtained from public stock centers solely as a limitation, multi-mutant designs enable these genetically complex populations to be leveraged for mapping and resolving genetic interactions.

We were able to map mutants in heterogeneous genetic backgrounds because of the genomic resources available capturing a large swathe of maize diversity [31,68]. Specifically, multiple reference genomes, as indispensable components of a pangenome [42], improved mapping resolution across diverse genetic backgrounds. Using multiple reference genomes provides a way to leverage genomic diversity and identify mapping intervals even for mutants with unknown or heterogeneous genetic backgrounds. High quality genome assemblies across species diversity are important even when mutagenesis is performed in known genetic backgrounds, because they can facilitate the mapping of natural variants and genetic modifiers [69–71]. As new model systems emerge [72,73], the development of high-quality genome assemblies and pangenomes will provide many opportunities for functional studies through forward genetics.

One major finding of this study is that most unmapped tasselseed mutants obtained from the Maize Genetics Cooperation Stock Center localized to genomic intervals containing known sex determination genes, including *Ts1*, *Ts2*, *Ts4*, *Ts6* and *Vrl1* [32,33,38–40,52]. Although these intervals do not establish allelism or identify causal genes, the recovery of mutants near these loci suggests that a relatively small number of developmental genes produce dramatic *tasselseed* phenotypes as single gene mutants. Collectively, these genes represent the functional roles of jasmonate metabolism (*Ts1* and *Ts2*), miRNA mediated regulation (*Ts4* and *Ts6*) and GT1/VRL1 transcriptional network in the modulation of carpel suppression and sex determination [32,33,37–40,52]. Notably, several mutants lacked obvious moderate or high impact variants in candidate genes within their mapping intervals. These findings suggest that some causal lesions may reside in noncoding regulatory regions that were not prioritized by our analysis. Alternatively, because the mutagenesis history of some unresolved mutants is unknown, they may result from variants other than canonical EMS SNPs that may have been missed by our filtering approach. In addition, because sequencing depth varied substantially among samples (6x–58x average genome coverage), some causal variants may have escaped detection in regions of limited coverage. Therefore, identifying underlying causal lesions of these mutants will require additional fine mapping and functional validation.

Our results demonstrate that forward genetic screens continue to uncover both putative new alleles of known sex determination genes and candidate novel loci. These mutants provide valuable genetic resources for dissecting the mechanisms underlying sex determination and carpel suppression in maize. More broadly, many phenotypic mutants preserved in public stock collections remain unmapped, representing an underutilized resource for gene discovery [30]. Continued mapping of these collections with next-generation genomics tools will facilitate the identification of additional alleles of known genes and novel loci regulating plant development.

## Materials and methods

### Plant materials and mapping populations

A total of 26 *tasselseed* mutants were obtained from the Maize Genetics Cooperation Stock Center (**S1 Table**) and crossed to various maize inbred lines to generate F2 mapping populations. F2 populations were grown, phenotyped, and analyzed for segregation ratios to determine the number of underlying loci. Following phenotyping, F2 individuals were pooled for bulked segregant analysis (BSA) using one of three experimental designs (mutant-only, wild-type versus mutant or multi-mutant). Equal amounts of leaf tissue were collected from each individual to construct a bulk. In the mutant-only design, only plants displaying strong tasselseed phenotype were pooled into a single pool (10-50 individuals). In the wild-type versus mutant and multi-mutant design, plants were grouped by phenotype, and equal amounts of leaf tissue were pooled from 16-84 individual plants into two or more bulks.

### DNA extraction and Next generation sequencing

Leaf tissue was collected as leaf discs using a single-hole puncher and pooled into a single 15 mL Falcon tube, as previously described by [19]. Each bulk contained 130-170 leaf discs, with an equal number of leaf discs taken from each individual plant to ensure equal representation and sufficient DNA yield for sequencing. Depending on the number of individuals selected for each bulk, 2 to 10 leaf discs were collected per plant. Leaf tissue was ground in liquid nitrogen and high molecular weight genomic DNA was extracted using a modified urea-based protocol [74]. DNA quantity and quality were assessed using Nanodrop and a Qubit fluorometer before library preparation. Sequencing libraries were prepared by Novogene Inc. and sequenced on an Illumina platform to generate 40 Gb of 150-bp paired-end reads per bulk.

### Read alignment and variant calling pipeline

De-multiplexed paired-end reads were downloaded from the Novogene server, and samples sequenced across multiple lanes were merged. Illumina adapter sequences and low-quality bases were trimmed using fastp [75] with default parameters and reads longer than 50 bp after trimming were retained for downstream analysis. Prior to alignment, each reference genome was indexed with minimap2, and reads were aligned with minimap2 using default parameters [76]. The resulting Sequence Alignment/Map (SAM) files were converted to Binary Alignment/Map (BAM) format, sorted and indexed using samtools. Read coverage and alignment statistics were computed using samtools flagstat and samtools depth [77]. Sorted and indexed BAM files were passed with bcftools mpileup (–ignore-RG) to generate genotype likelihoods and variants were called using bcftools call (-m and -v options) [77]. Variant Call Format (VCF) files were converted to a tab-delimited table using bcftools query. The resulting table included the following columns: chromosome (CHROM), position (POS), reference allele (REF), alternate allele (ALT), variant quality score (QUAL), total read depth (DP), and strand-specific read counts extracted the DP4 field, renamed as forward reference (Fref), reverse reference (Rref), forward alternate (Falt), and reverse alternate (Ralt) for clarity.

### Allele frequency calculation and BSA statistical analysis

Following read mapping and variant calling, the downstream analyses such allele frequency calculation and visualization were performed using R [78] in the RStudio environment [79] using customized R scripts. The tab-limited variant tables were imported into R, and only single nucleotide polymorphisms (SNPs) were retained whereas the insertions/deletions (indels) were excluded. SNPs were further filtered to include only those SNPs with a minimum read depth (DP) of 5-15 and minimum variant quality score (QUAL) of 10-30 depending on sequencing coverage.

For each retained SNP, alternate allele frequencies were calculated using strand-specific read counts as the sum of alternate reads (Falt and Ralt) divided by the total reads at that SNP position (Falt, Ralt, Fref and Rref). These allele frequencies (AF) were used to calculate allele frequency differences (AFD), homozygosity mapping (HM), G-statistic (G), Euclidean distance, (ED) and its fourth power transformation (ED^4^). AFD was calculated as the difference in alternate allele frequencies between bulks [45]. G-statistic (G) was calculated using the observed and expected allele counts from the two bulks as described by [46]. In contrast, ED and ED^4^ were calculated using both reference and alternate allele frequencies, where reference AF was computed as one minus the alternate AF. ED was calculated as the square root of the sum of squared differences in alternate and reference AFs between bulks [80], and ED⁴ was obtained by raising the ED value to the fourth power [47]. Genome-wide variations were visualized with plots generated using the ggplot2 package in R [81] . However, plots of raw AFs, AFD, ED and ED^4^ were often noisy, local regression smoothing was applied using a locfit package in R. A local polynomial regression curve was fitted using the nearest 10% of SNPs surrounding each genomic position [82].

As an additional analysis to identify candidate mapping intervals, genome-wide values of AFD, G, ED^4^, and HM were summarized using a sliding window approach of a 2 Mb window and a 100 kb step size. SNPs with a minimum read depth of 15 and a minimum variant quality of 20 were retained for sliding window based analysis. In each window, the median AFD, G, and ED^4^ values were calculated, whereas HM was calculated as the proportion of homozygous SNPs (AF ≥ 0.99 or AF ≤ 0.01). Optional rolling median and local polynomial (locfit) smoothing were applied for visualization. To identify genomic regions enriched for mapping signal, a sliding-window Fisher’s exact test was performed for HM, AFD and ED^4^. For HM, AFD, G and ED^4^, SNPs were classified as above or below threshold of 0.99, 10, 0.75 and 1.5, respectively, and Fisher’s exact test compared the number of SNPs above the threshold within each window to the remainder of the genome [82]. P-values were adjusted for multiple testing using the Bonferroni correction and reported as –log_10_(adjusted P-values) (**S5 Fig)**.

### Structural variant calling and Candidate SNP identification

Sorted and indexed BAM files were analyzed with Manta to identify structural variants (SVs) from next-generation sequencing data [83]. Manta was run with default parameters to identify a range of SV types, including insertions, deletions, inversions and duplications, by leveraging discordant read pairs and split-reads to detect structural variation. To refine the mapping interval of the mutant of interest, structural variants within the candidate region were identified and custom molecular markers were designed based on these variants to speed up fine mapping.

SVs especially insertions and deletions of 100 base pairs or larger were selected. and Primers flanking each variant by 100-200 base pairs were designed using primer3 [84] to enable PCR and gel-based genotyping. DNA for individual plants was extracted, and genotyping was performed as described by [85].

The effects of SNPs on gene function were predicted and annotated using SnpEff [48] relative to the B73 reference genome. Within each mapping interval, candidate causal variants were prioritized based on the expected mutational signature of the mutagen. For EMS induced mutants, only canonical G/C to A/T transitions were retained after excluding background SNPs present in the maize Hapmap VCF [86]. Candidate variants were further prioritized according to their predicted functional impact, with high and moderate impact variants given the highest priority for downstream analysis.

## Data availability

All raw sequencing reads are being deposited in the NCBI Sequence Read Archive (SRA) database under accession number PRJNA1505212. The BSA-Seq bioinformatic pipeline is available on GitHub at https://github.com/bzebosi/BSA-Seq. Seeds of the mutants used in the study are available at the Maize Genetics Cooperation Stock Center.

## Author contributions

M.B. and S.L. conceptualized the research, acquired funding and supervised the project. B.Z. and C.B.A. performed the experiments. B.Z. curated and analyzed the data. B.Z. wrote the original draft of the manuscript. B.Z., M.B., and S.L., revised and edited the manuscript. All authors read and approved the manuscript.

## Acknowledgments

The authors thank Maize Genetics Cooperation Stock Center staff, Marty Sachs and Jeff Gustin for providing seed stocks and valuable feedback. Oregon State University, Botany and Vegetable research farm staff for nursery efforts especially field preparation and irrigation. This work was supported by NSF-PGRP grant (IOS-2211434) and USDA NIFA grant (2023-67013-44037).

## Supporting information

**S1 Fig. Comparison of BSA-Seq mapping statistics for the 6609A (*ts*04HI*) mutant** Smoothed genome-wide mapping plots of AFD, ED, G and ED^4^ for the 6609A (*ts*04HI*) mutant with reads aligned to the Mo17 reference genome. Although all four methods identified a mapping peak on chromosome 2, G and ED^4^ produced narrower peaks, with ED^4^ showing reduced background noise. The red shaded region indicates the mapping interval, and the arrow marks the location of *ts1* locus.

**S2 Fig. Smoothed mapping plots of mutants localized to chromosomes 1, 3, 4 and 7. A-D)** Chromosome-specific BSA-Seq plots. The y-axis shows AF and ED4 mutants mapped using mutant-only and wild-type versus mutant designs, respectively, and the x-axis shows chromosome position (Mb). B73 was used as the reference genome for all mutants except *6609B* and *6609I* (Mo17) and 5806L (P39). **A)** Chromosome 1 intervals included *Ts2* (*4012U*, *6508F*, *6508P*, *6609B*, *6508C*, *mb7 ts*-07IL*, and *6609I*), Ts6 *(4012R*), or a putative locus (*tts*-2481)*. **B)** Chromosome 3 interval of *5807K* contained *Ts4* (∼146.6 Mb). **C)** No known *tasselseed* genes were identified within chromosome 4 interval of *4012M.* **D).** The chromosome 7 interval of *5806L* overlapped *Vrl1* (∼169.9 Mb). Brown shaded regions indicate mapping intervals, and arrow mark known tasselseed loci

**S3 Fig. Smoothed chromosome 2 mapping plots.** All mutants mapped to intervals containing the *Ts1* gene (∼47.9 Mb, arrow). B73 was used as the reference genome, except for *6508B* (P39), *6711D* and *ts*-03IL* (W22). Brown shaded areas indicate mapping intervals. The y-axis shows ED⁴, and the x-axis shows chromosome position (Mb).

**S4 Fig. Smoothed plots for mutants mapped on chromosome 2. A-C)** Mapping plots for mutants whose intervals were difficult to resolve using genome-wide ED4 alone but were refined using Fisher’s exact test. Upper panels show smoothed ED4 values, and the lower panels show sliding-window –log10(adjusted P-values) from Fisher’s exact test using a 2 Mb window and 100 kb step size. Mo17 was used as a reference genome for read alignment. The shaded region and arrow indicate mapping intervals and arrows mark the *Ts1* locus (∼47.9 Mb).

**S5 Fig. *4012M* (*ts*-N1967A*) localizes to a candidate interval on chromosome 4. A-B)** Mapping of *4012M* (*ts*-N1967A)* using two independent mapping populations in the B73 (**A**) and W22 (**B**) genetic backgrounds, with reads aligned to the P39 reference genome. The upper panels show genome-wide ED4 values smoothed using locfit with a nearest neighbor proportion of 0.1 (nn = 0.1). The lower panels show sliding-window –log10(adjusted P-values) from Fisher’s exact test using a 2 Mb window with a 100 kb step size. Variants with a minimum read depth of 15 and quality score of 20 were used for Fisher’s exact test, which compared the number of SNPs with ED^4^ >= 1.5 and ED^4^ < 1.5 between each window and the remainder of the genome. The black horizontal dotted line indicates the significance threshold (adjusted P < 0.001), and shaded region indicates mapping interval.

**S1 Table**. Tasselseed mutant stocks used for the study

**S2 Table.** Summary of sequencing and alignment statistics for samples.

**S3 Table.** Read alignment statistics for samples across reference genomes.

**S4 Table.** Mapping intervals and candidate genetic variants in tasselseed mutants.

