## supplemental figures for "BSA101: Unlocking Historical Mutant Collections with BSA-Seq"

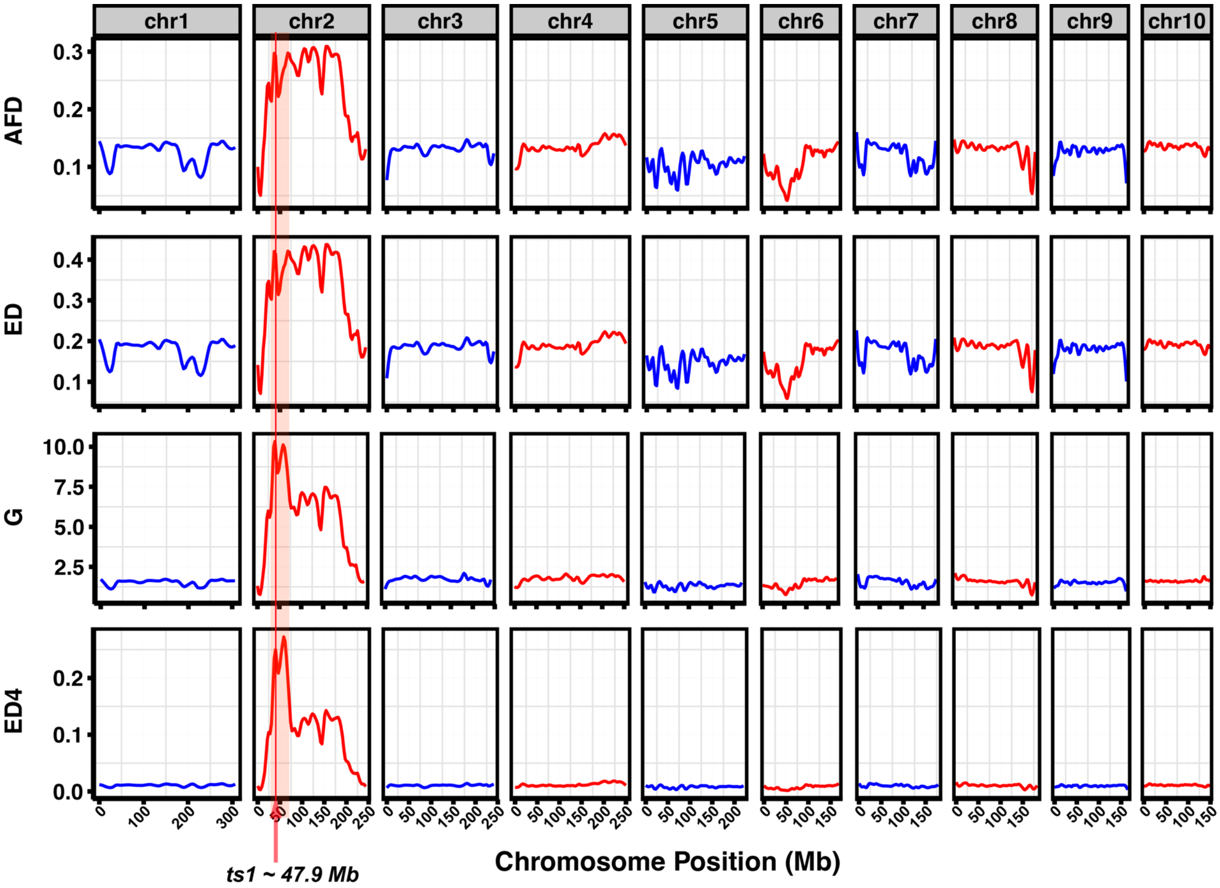


**S1 Fig. Comparison of BSA-Seq mapping statistics for the 6609A (*ts*04HI*) mutant**

Smoothed genome-wide mapping plots of AFD, ED, G and $ED^4$ for the 6609A (*ts*04HI*) mutant with reads aligned to the Mo17 reference genome. Although all four methods identified a mapping peak on chromosome 2, G and $ED^4$ produced narrower peaks, with $ED^4$ showing reduced background noise. The red shaded region indicates the mapping interval, and the arrow marks the location of *ts1* locus.

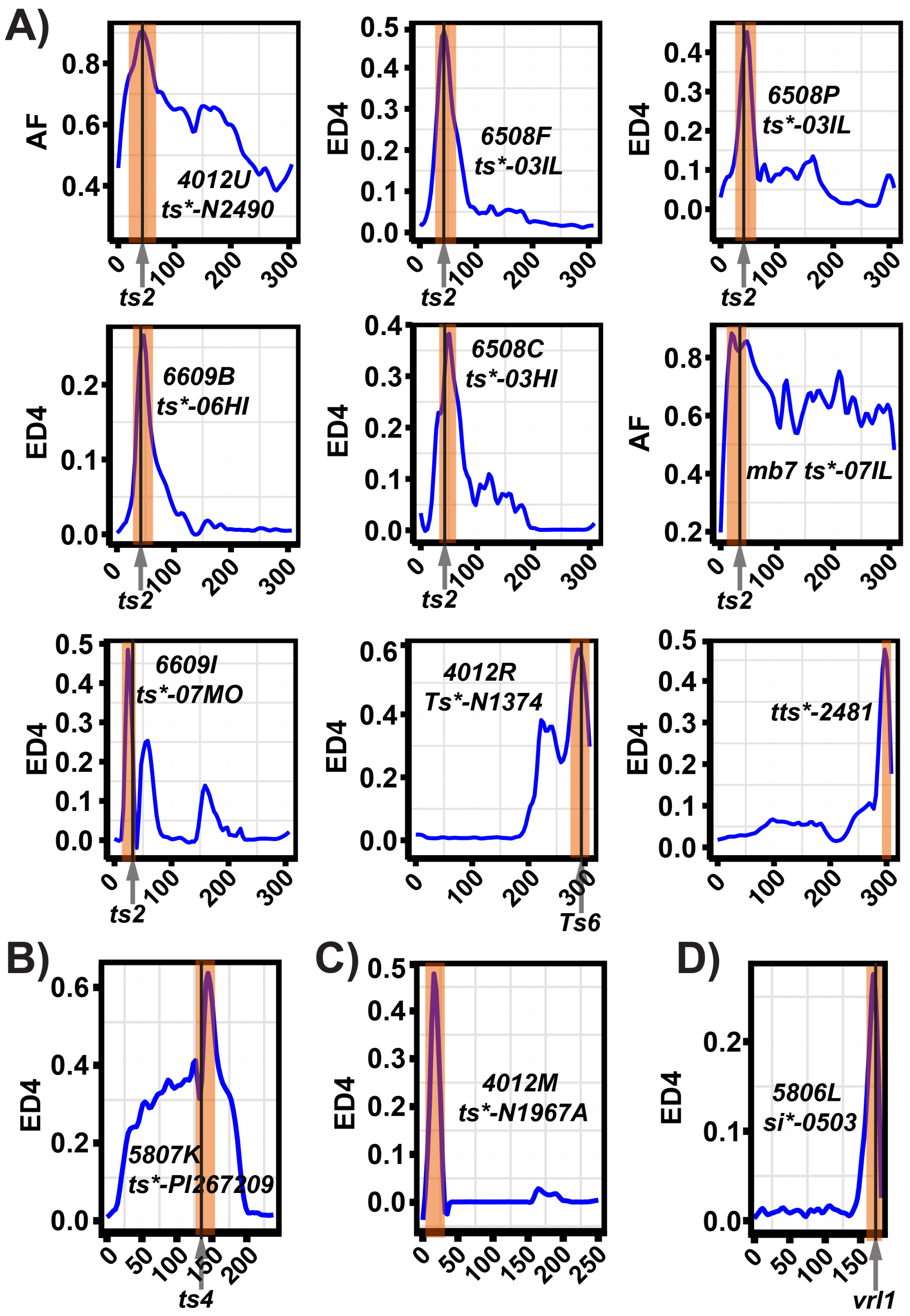


**S2 Fig. Smoothed mapping plots of mutants localized to chromosomes 1, 3, 4 and 7.**
**A-D)** Chromosome-specific BSA-Seq plots. The y-axis shows AF and ED4 mutants mapped using mutant-only and wild-type versus mutant designs, respectively, and the x-axis shows chromosome position (Mb). B73 was used as the reference genome for all mutants except *6609B* and *6609I* (Mo17) and 5806L (P39). **A)** Chromosome 1 intervals included *ts1* (*4012U*, *6508F*, *6508P*, *6609B*, *6508C*, *mb7 ts*-07IL*, and *6609I*), *Ts6 (4012R*), or a putative locus (*tts*-2481)*. **B)** Chromosome 3 interval of *5807K* contained *Ts4* (~146.6 Mb). **C).** No known *tasselseed* genes were identified within chromosome 4 interval of *4012M.* **D).** The chromosome 7 interval of *5806L* overlapped *Vrl1* (~169.9 Mb). Brown shaded regions indicate mapping intervals, and arrow mark known tasselseed loci

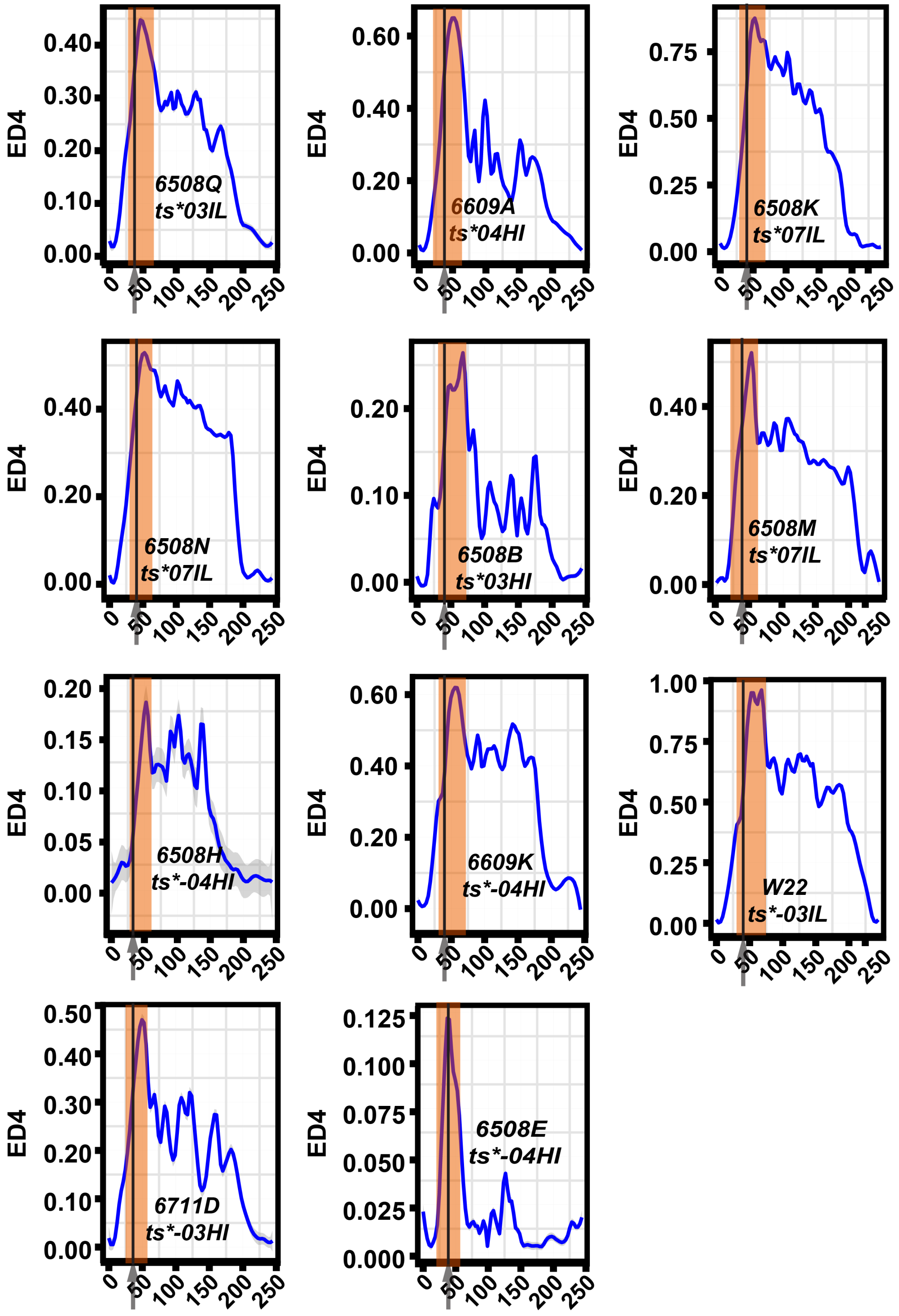


**S3 Fig. Smoothed chromosome 2 mapping plots.** All mutants mapped to intervals containing the *Ts1* gene (~47.9 Mb, arrow). B73 was used as the reference genome, except for *6508B* (P39), *6711D* and *ts*-03IL* (W22). The y-axis shows $ED^4$, and the x-axis shows chromosome position (Mb). Brown shaded areas indicate mapping intervals and arrow mark *Ts1* locus.

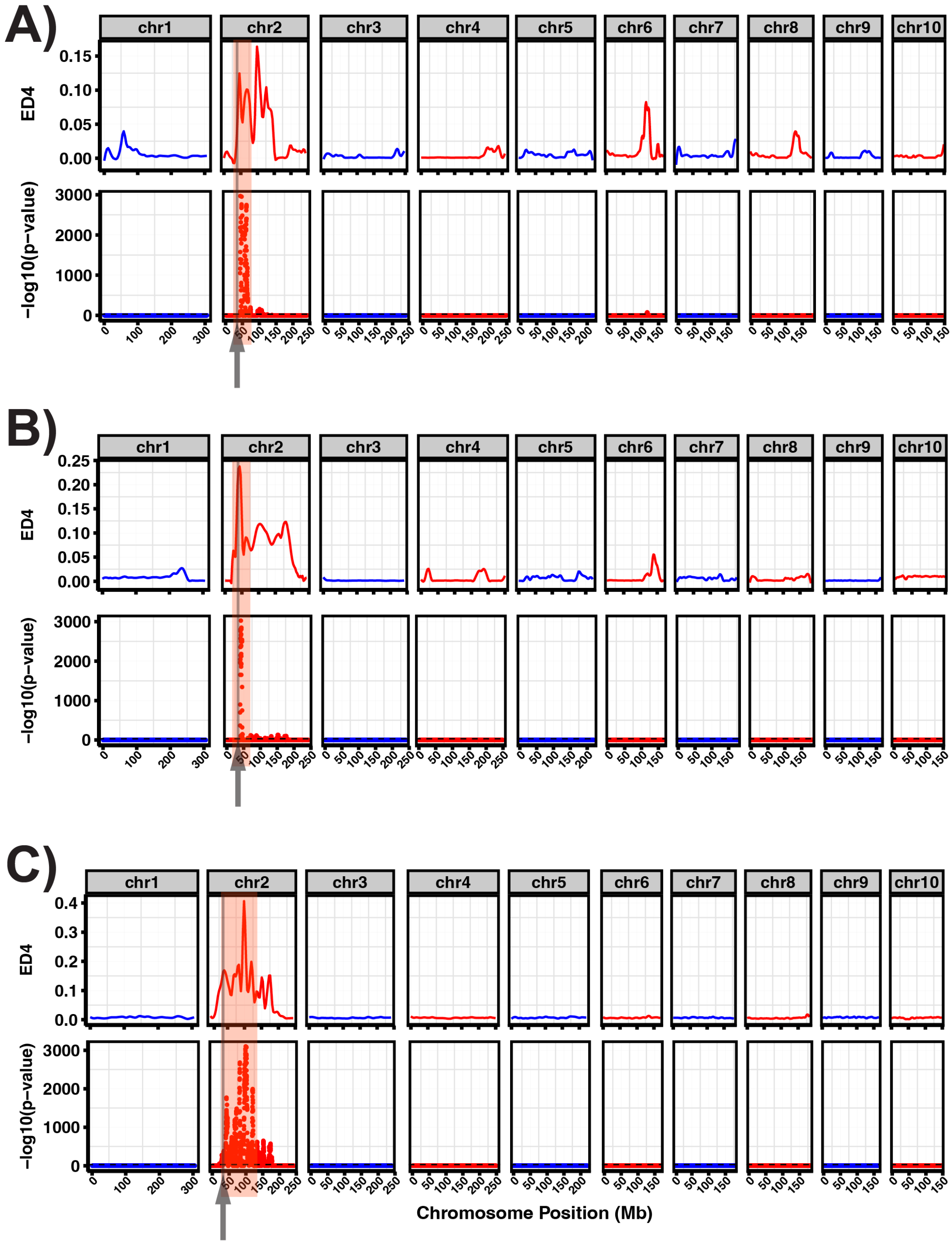


**S4 Fig. Smoothed plots for mutants mapped on chromosome 2.**
**A-C)** Mapping plots for mutants whose intervals were difficult to resolve using genome-wide ED4 alone but were refined using Fisher's exact test. Upper panels show smoothed ED4 values, and the lower panels show sliding-window –log10(adjusted P-values) from Fisher's exact test using a 2 Mb window and 100 kb step size. Mo17 was used as reference genome for read alignment. Red shaded region and arrow indicate mapping intervals and arrows mark the *Ts1* locus (~47.9 Mb).

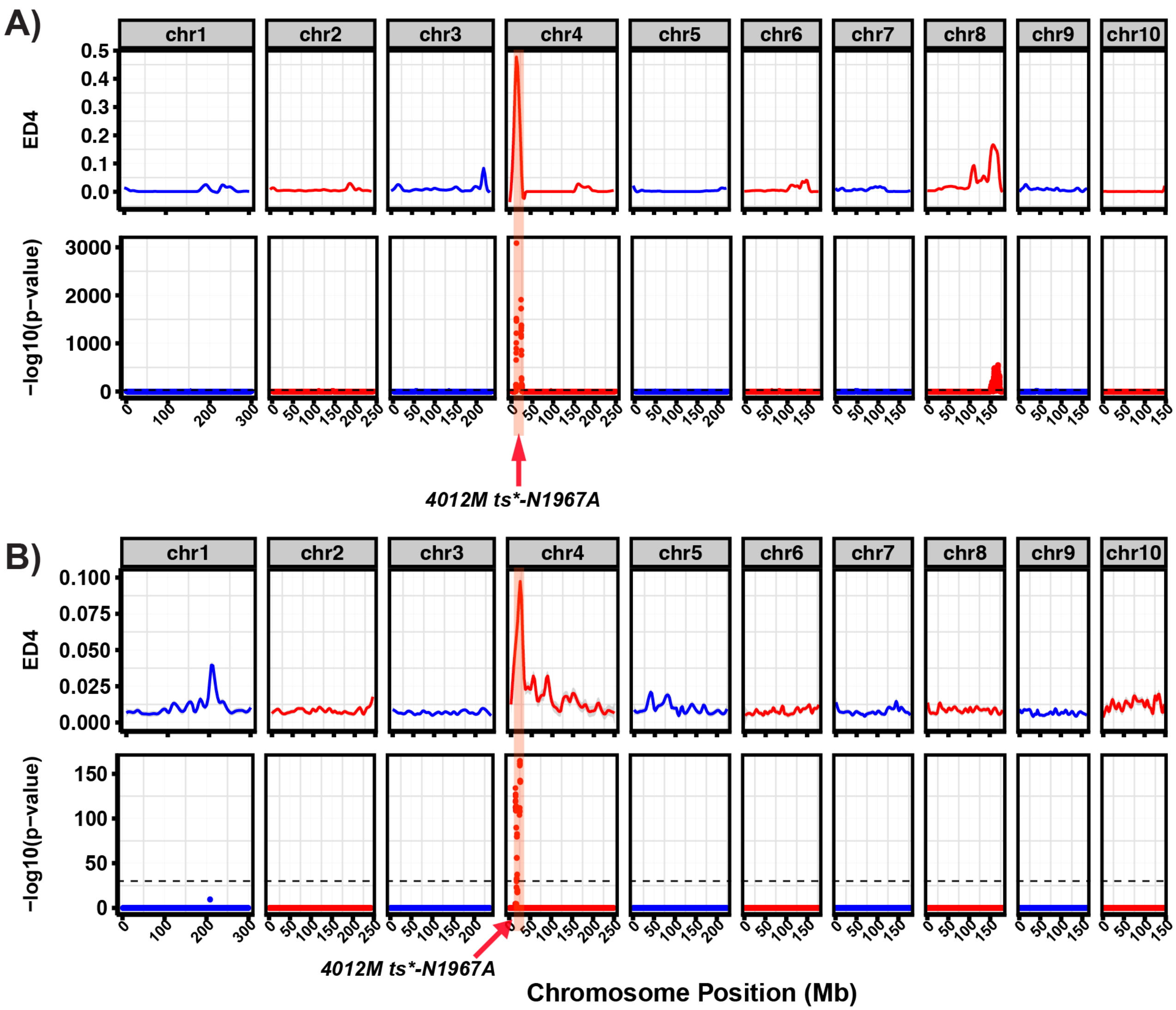


**S5 Fig. *4012M* (*ts*-N1967A*) localizes to a candidate interval on chromosome 4.**
**A-B)** Mapping of *4012M* (*ts*-N1967A)* using two independent mapping populations in the B73 (**A**) and W22 (**B**) genetic backgrounds, with reads aligned to the P39 reference genome. The upper panels show genome-wide ED4 values smoothed using locfit with a nearest neighbor proportion of 0.1 (nn = 0.1). The lower panels show sliding-window –log10(adjusted P-values) from Fisher's exact test using a 2 Mb window with a 100 kb step size. Variants with a minimum read depth of 15 and quality score of 20 were used for Fisher's exact test, which compared the number of SNPs with ED4 >= 1.5 and ED4 < 1.5 between each window and the remainder of the genome. The black horizontal dotted line indicates the significance threshold (adjusted P < 0.001), and shaded region indicates mapping interval.
